# The Role of Purine Interactions in Biogenic Crystal Shape Determination

**DOI:** 10.1101/2024.09.18.613275

**Authors:** Jannik Rothkegel, Sylvia Kaufmann, Michaela Wilsch-Bräuninger, Rita Mateus

## Abstract

Widespread through phyla, purine crystals are intracellular inclusions serving a myriad of organismal functions. In zebrafish, iridophores concentrate purines in membrane-bound organelles, the iridosomes, for controlled crystallization. These crystals assemble into large, flat, and thin hexagons following unknown mechanisms that evolve against thermodynamically favorable interactions. Here, we investigate the initial development of zebrafish iridosomal crystals. By performing *in vivo* confocal reflection imaging, cryoFIB-SEM, and establishing novel 2D and 3D analysis pipelines, we show that these crystals grow four times faster along the b-crystallographic axis, leading to their characteristic hexagonal shape. By analyzing zebrafish with impaired guanine production, while conducting crystal growth simulations, we find that crystal shape is directed by bond type, number, and interaction strength between purines. Mechanistically, the macroscopic shape of zebrafish crystals is controlled by the relative concentration of purines present in the iridosome. This process impacts crystal growth along the b-axis, by disrupting the crystal’s in-plane hydrogen bond structure, without alteration of the other axes. Our work uncovers a layer of biogenic crystal growth regulation occurring in vertebrate biocrystallization processes.

**Highlights:**

- Zebrafish iridophores actively regulate crystal size and shape within iridosomes
- *In vivo* crystal (100) facet grows ∼4μm² in 24 hours
- Length of crystallographic b-axis is controlled by purine molecular interactions
- Size of c- and a-axes is regulated independently of b-axis

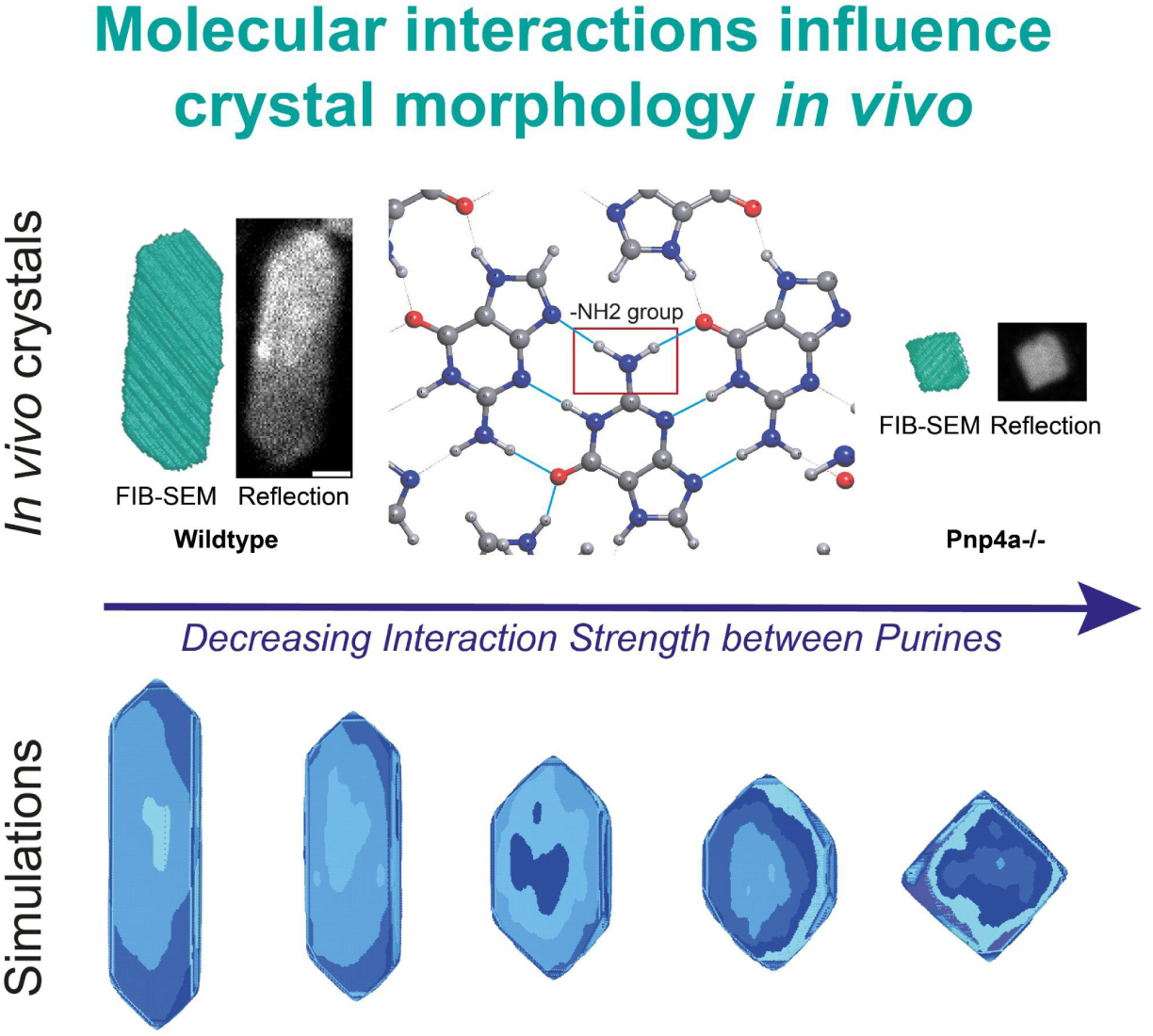

## Introduction

Many organisms utilize organic crystals made of purines, the most abundant of which is the nucleobase guanine^1–3^, to manipulate light^4^. They are used for a wide range of functions, including the production of structural colors^5–7^, heat protection^8^, and even optical components in the visual system^9,10^. Purine crystals can fulfill this plethora of tasks as, depending on their size, morphology, and structural arrangement, they can be used as light scatterers, multilayer reflectors, or image-forming mirrors^4^. The very high refractive index of guanine crystals (n=1.83) along the pi-pi stacking direction^11^ and the fact that guanine is readily available as an end-product of nucleic acid degradation^12^, makes these molecules ideal for building optical elements inside animals efficiently. As crystal morphology is tightly coupled to their intended cellular function, organisms are capable of robustly influencing and controlling this biocrystallization process. Importantly, disruption of crystal biogenesis undermines organismal fitness and can be detrimental to tissue performance, and can lead to pathologies^13,14^.

In vertebrates, purine crystals are produced by specialized pigment cells called iridophores or leucophores, which differentiate from the neural crest^15,16^. These crystals adopt a beta-polymorph structure^17^, composed of hydrogen-bonded layers, which are stacked via pi-pi interactions, leading to diverse prismatic shapes^18^. In zebrafish (*Danio rerio*), each crystal is intracellularly encapsulated in a membrane-bound organelle - the iridosome^19–21^. In this vesicle, crystallization is thought to occur via the accumulation of supersaturated guanine, which undergoes phase transitions into amorphous and then crystalline phases^22^. Recent studies have revealed that biogenic organic crystals, including those of zebrafish, are not only composed of pure guanine, but also include varying amounts of other purines, including up to 18% hypoxanthine^2^. Further, by genetically disrupting purine nucleoside phosphorylase 4a, i.e. Pnp4a, a crucial teleost protein in the purine metabolic network of iridophores^23–25^, we have recently shown that reducing guanine production leads to smaller, differently-shaped crystals, resulting from altered concentration ratios of hypoxanthine-to-guanine^23^. However, precise quantification of these morphological changes and a detailed understanding of the molecular interactions driving this crystal shape change remain unexplored.

Herein, we use confocal imaging and electron microscopy techniques to track crystal growth in early zebrafish development from 30 to 96 hours post-fertilization (hpf). We demonstrate that biogenic zebrafish crystals preferentially grow along the b-axis, giving rise to their typical elongated shape. By quantitatively comparing crystal morphometrics between wildtype and Pnp4a-/- mutants, we show that the latter crystals, despite their square morphologies due to a severely shortened b-axis, have unaltered thickness and width when compared to wildtype. By performing molecular-level Monte Carlo simulations, we identify that the observed Pnp4a-/- crystal morphologies are caused by a disruption of the crystal hydrogen bond network. This mechanistic framework can be generalized to explain how crystallographic axis length is regulated in biogenic organic crystals via molecular interactions.

## Results

### Morphometric characterization of crystal size and shape in zebrafish iridophores

To elucidate the iridosomal crystal morphology as they develop inside differentiating iridophores, we performed live imaging of transgenic reporter zebrafish that express GFP specifically in iridophores, Et(Ola.Edar:GAL4,14xUAS:GFP)^TDL358Et^ (i.e. TDL358:GFP from here onwards)^26^. We decided to use the larval eye as our developmental model, as this is the first organ that contains iridophores that develop iridosomes, enclosing crystals with stereotypic shapes and sizes beyond the diffraction limit^27^. Given the orientation of these vesicular crystals within the cytosol of the iridophores, the planar exposed (100) crystal facets reflect incident light when illuminated, which we took advantage of by using point-scanner confocal microscopy, in both fluorescent and reflection modes (Fig. 1A-C). By quantifying crystal reflection within iridophores and the respective surface area covered by iridophores in the eye across time, we found that total eye reflection increases exponentially during iridophore development (Fig.1E). In contrast, the total iridophore area covering the eye increases linearly in time (Fig.1D). Together, this indicates that during this developmental period (30-96 hpf), the iridophores actively produce large amounts of intracellular crystals, leading to the dense intracellular packing observed with iridosomal guanine crystals (Fig.1C-E).

**Figure 1.**
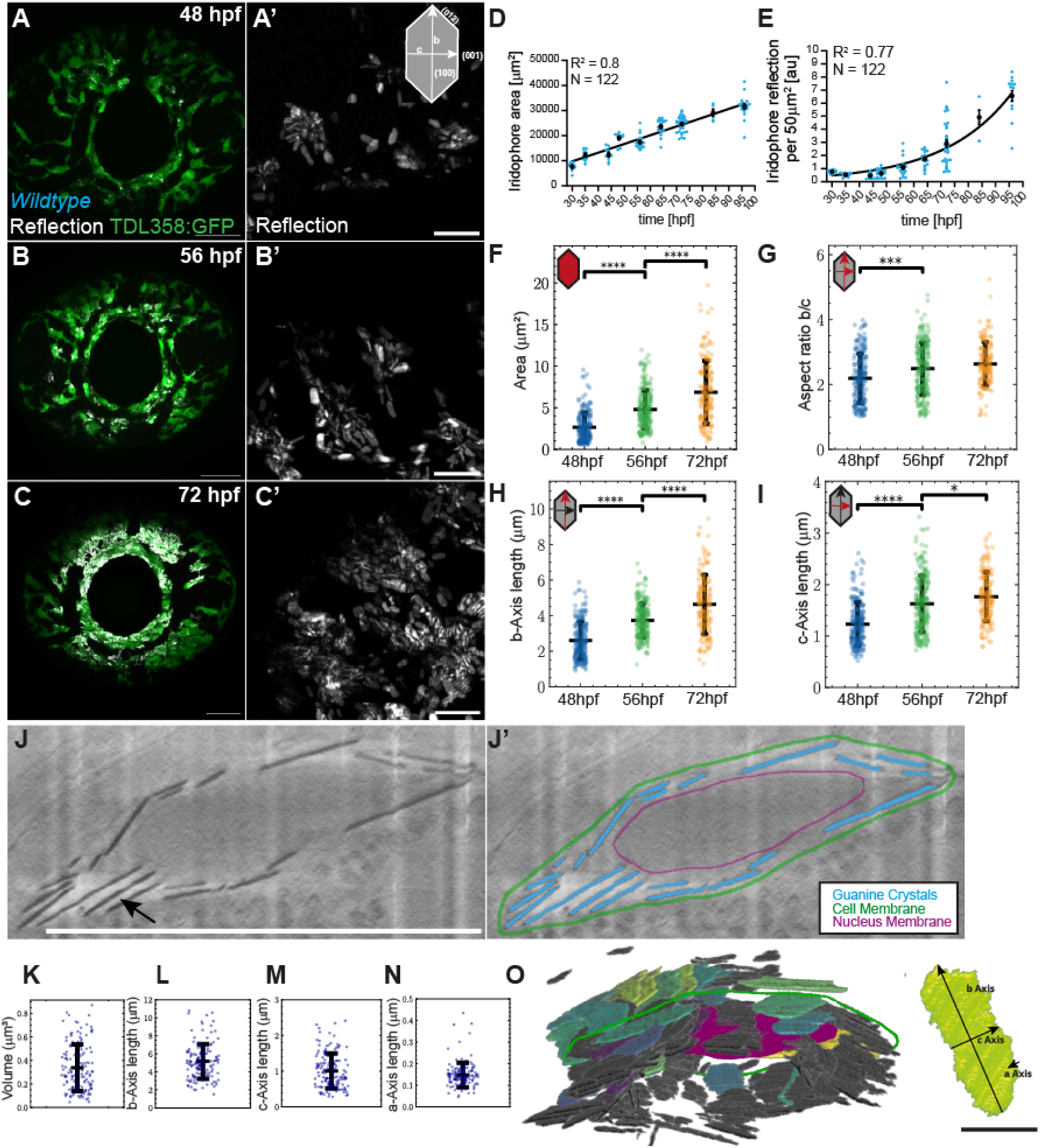
Crystal reflection onset and morphology in larval zebrafish. **A-C** Confocal microscopy maximum intensity projection images of larvae eyes of iridophore reporter line (TDL358:GFP, green) and reflection (white), at 48 hpf (**A**), 56 hpf (**B**) and 72 hpf (**C**), with zoom in on crystals from the same time points (**A’-C’**). Cartoon indicates crystallographic axes in 2D with respective crystal facets. Scale bars: A-C 50μm, A’-C’ 10μm. **D-E** Iridophore area in the eye increases linearly between 30 and 96 hpf (**D**; Line, linear fit with goodness of fit, R^2^), while iridophore reflection increases exponentially within the same time interval, in the same cells (**E,** Line, exponential fit with goodness of fit, R^2^). Mean ± SEM are shown in black. **F-I** Morphometric measurements from 2D segmented crystals obtained from reflection imaging of wt zebrafish eyes at three different time points. Area (**F**), Aspect Ratio of b-axis/c-axis Lengths (**G**), b-axis Length (**H**), and c-axis Length (**I**). Asterisks indicate p-values of unpaired, two-tailed, non-parametric t-tests between groups at each time point: ∗p<0.05, ∗∗∗p<0.001, ∗∗∗∗p<0.0001. Error bars, standard deviation of the mean. N, number of crystals. 48 hpf, N=197; 56 hpf, N=227; 72 hpf, N=107. **J** CryoFIB-SEM slice of wild-type iridophore at 72 hpf, with respective segmentation (**J’**). Guanine crystals appear dark due to their comparatively high density (black arrow). **K-N** Measurements from 3D segmented crystals obtained from cryoFIB-SEM imaging; N=139 crystals. Volume (**K**), b-axis length (**L**), c-axis length (**M**), and a-axis length (**N**). **O** 3D reconstruction from J. Fully segmented crystals are colored relative to volume, with darker colors corresponding to lower volume. Cell membrane (green) and nucleus (magenta) labels correspond to the ones depicted in K. On the right, a single reconstructed crystal with crystallographic axes marked. Scale bars, 10μm (J), 1μm (O).

To extract 2D morphometric features considering the crystal’s reflective (100) facet, we developed and implemented a 2D segmentation pipeline that allowed us to quantify large numbers of individual crystals *in vivo* (see Methods). We found that, at 48 hpf, the crystals have an area of approximately 2.63μm², which increases to 4.78μm² at 56 hpf and 6.84μm² at 72 hpf (Fig. 1F). Interestingly, the contribution of the two crystal axes (b and c) to this increase in area is distinct. The length of the b-axis measures 2.60μm at 48 hpf and 4.63μm at 72 hpf, while the c-axis length increases from 1.23μm to 1.77μm, showing that the b-axis grows four times faster than the c-axis (Fig. 1H,I). Interestingly, the crystals’ aspect ratio (b/c) does not significantly change after 56 hpf, indicating that crystal proportions are established from this time point onwards (Fig. 1G). Additionally, we find that there is always a population of small crystals (< 2μm²) (Fig. S1), indicating that crystal growth is heterogeneous and not fully synchronized per cell. Rather, crystal growth starts individually inside each iridosome, most likely depending on the iridosomal physico-chemical conditions and the cell’s differentiation state^15,16^.

To increase the accuracy and resolution of our morphometric measurements, we conducted cryogenic Focused Ion Beam Scanning Electron Microscopy (cryoFIB-SEM) of zebrafish eyes, specifically acquiring images of the iridophores at 72 hpf (Fig. 1J). This volumetric technique preserves the native conditions of tissues and cells without destroying crystal vesicles or the crystals themselves^28^. Importantly, this approach also allowed us to characterize the crystals’ a-axis (i.e. thickness) quantitatively, given this axis’ nanometer length scale^29^. By establishing a semi-automated 3D segmentation and reconstruction image analysis pipeline, we extracted morphometric measurements of all three crystallographic axes, with respective volumes (see Methods). This revealed that crystals have the expected hexagonal morphology, independently of the current size (Fig. 1L-M,O). The crystals display an average volume of 0.33μm³ (Fig. 1K), with b- and c-axis lengths with averages of 5.2μm and 1μm for each axis, respectively (Fig. 1 L,M). By comparing with our live imaging data, we verified that those measurements are within the standard deviation range of our cryoFIB-SEM data, which captures such axes in greater detail (compare Figs. 1H,I with L,M). The a-axis (∼140nm on average) has a narrow length distribution compared to the other axes (Fig. 1N), indicating that thickness is tightly regulated within iridosomes.

Taken together, we conclude that when compared to other crystal facets, the (012) facet has a higher growth rate, promoting the expression of the (100) and (001) facets. Because of this, the guanine crystals obtain their characteristic hexagonal elongated shape.

### The macroscopic shape of zebrafish crystals is controlled by the relative concentration of purines in the iridosome

Recent studies from our lab and colleagues show that the purine composition is crucial to defining crystal morphology and, consequently, its function^23,30^. Given that zebrafish crystals are homogeneous solid solutions composed of not only guanine, but also other purines, namely hypoxanthine^2,23^, we hypothesized that the molecular interactions between the crystal atoms, such as H-bonds, significantly affect crystal growth modes and respective growth directions.

In line with the above, we have previously shown that mutating the purine nucleoside phosphorylase 4a (*pnp4a*) gene in zebrafish effectively leads to a decrease in guanine concentration, as this protein catalyzes the cleavage of guanosine/deoxyguanosine into guanine^23,25^. *Pnp4a^cbg^*^20^ homozygous mutants (Pnp4a-/-) were still able to produce crystals, albeit in small amounts, which surprisingly appeared to show a rhomboid-like morphology^23^. By performing confocal live imaging of reflection and applying our quantitative image analysis pipelines, we first confirmed that *pnp4a* is highly expressed in iridophores (Fig. 2A)^31^. We then expanded our quantification efforts to compare Pnp4a-/- crystal morphologies with those of wildtype zebrafish. We observed that crystals produced by Pnp4a-/- mutants exhibit distinct square shapes (Fig. 2B) with an aspect ratio close to 1 (Fig. 2D). The mutant crystals are smaller compared to wildtype, with an average area of 2.2μm² compared to 4.78μm² at 56 hpf, respectively. At 72 hpf, Pnp4a-/- crystals do not grow beyond 2.2um² on average per crystal, while in the wildtype case, those reach an average of 6.84μm² per crystal (Fig. 2C, S1). The lack of growth of the b-axis (average of 1.7μm at 56 and 72 hpf) results in the Pnp4a-/- crystals’ phenotype, presenting underdeveloped (001) crystal facets, further validating diffraction and *in vitro* findings^23^.

**Figure 2.**
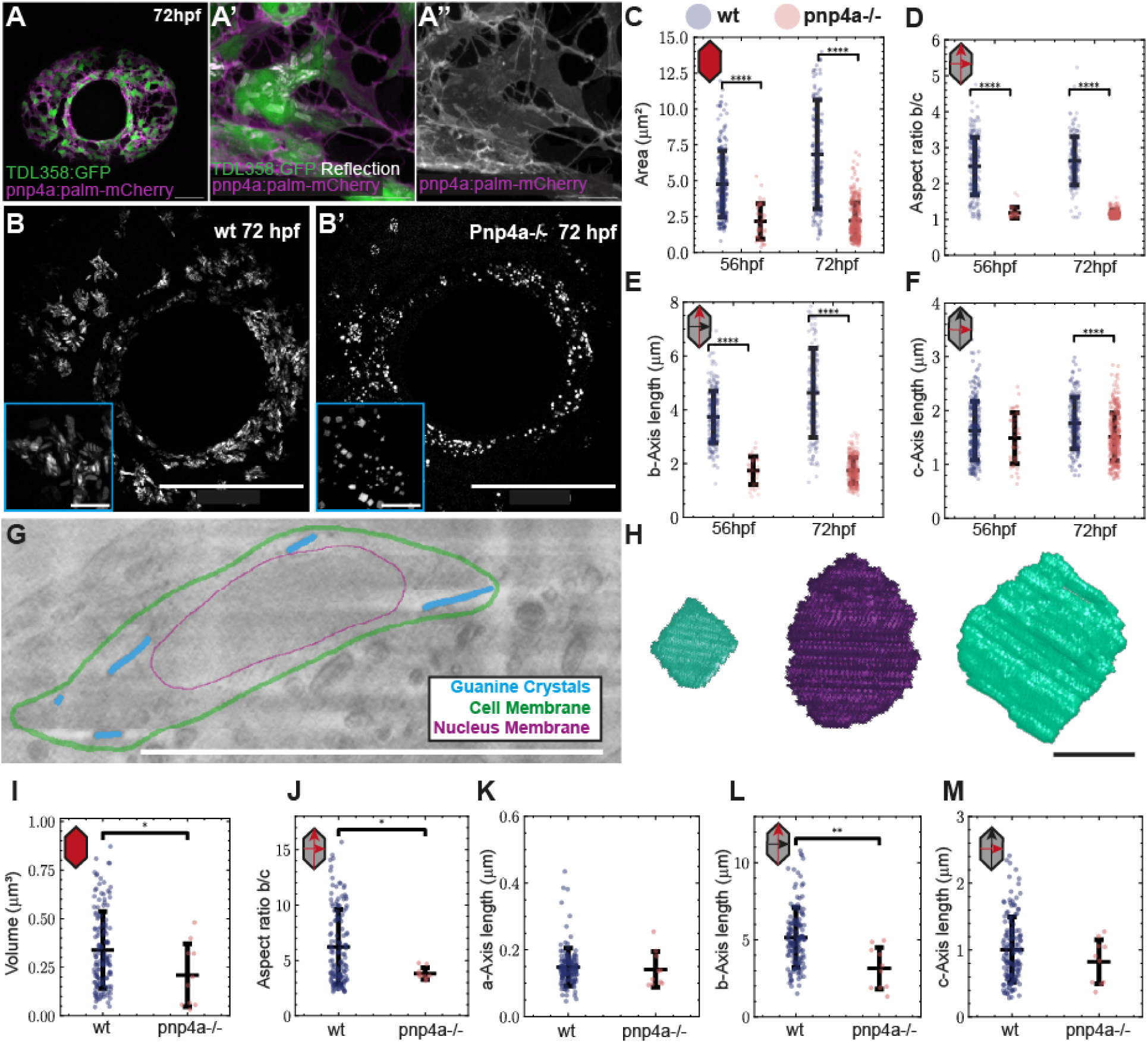
Pnp4a-/- crystals display distinct square morphologies due to an underdeveloped (001) crystal face. **A** Eye iridophores labeled in double transgenic TDL358:GFP (green) and pnp4a:PALM-mCherry (magenta), with zoom highlighting the co-localization of expression as well as reflection signal, showing respective crystals (white), at 72hpf (**A’-A’’**). Scale bars: 50μm (A), 10μm (A’-A’’). **B-B’** Confocal reflection images of larval eyes in wildtype (**B**) and Pnp4a-/- (**B’**) at 72 hpf. Blue zoom insets highlight respective crystal morphologies. Scale bars: 100μm, 10μm (inset). **C-F** Comparisons of 2D morphometric crystal measurements between wildtype (blue) and Pnp4a-/- mutants (red) at 56 and 72 hpf. Area (**C**), Aspect Ratio of b-axis/c-axis Lengths (**D**), b-axis Length (**E**), and c-axis Length (**F**). N, number of crystals. 56hpf, WT N=227, Pnp4a-/- N=41; 72hpf, WT N=107, Pnp4a-/- N=315. **G** CryoFIB-SEM slice with segmented iridophore from Pnp4a-/- mutants at 72 hpf. **H** cryoFIB-SEM reconstructed crystals from Pnp4a-/- zebrafish express square morphologies. Scale bars: 10μm (G), 1μm (H). **I-M** Comparison of 3D morphometric measurements between wildtype (blue) (N=139) and Pnp4a-/- crystals (red) (N=10). Volume (**I**), Aspect Ratio of b-axis/c-axis Lengths (**J**), a-axis Length (**K**), b-axis Length (**L**), and c-axis Length (**M**). Asterisks indicate p-values of unpaired, two-tailed, non-parametric t-tests between groups: ∗p<0.05, ∗∗p<0.01, ∗∗∗∗p<0.0001. Error bars, standard deviation of the mean.

To study crystal morphology in the Pnp4a-/- mutants in more detail, we again conducted cryoFIB-SEM in zebrafish eye iridophores at 72 hpf. 3D segmentation, reconstruction, and quantification of the acquired cryoFIB-SEM images, confirmed our initial confocal observations that the iridophores in Pnp4a-/- mutants produce fewer iridosomes filled with crystals, these being very interspersed within the iridophore cytosol (Fig. 2G)^23^. The volume of Pnp4a-/- crystals was lower than that of wildtype crystals, at an average of 0.21μm³ (Fig. 2I). In line with our light microscopy data and previous work^23^, we found that the largest contributor to this volume change is indeed the shorter b-axis, with an average length of 3.16μm in Pnp4a-/- mutants compared to 5.15μm in wildtype (Fig. 2L,M). Interestingly, the lengths of the a- and c-axes did not change significantly (Fig. 2K,M). This provides quantitative evidence supporting the notion of a reduced growth rate of the (012) facet in Pnp4a-/- mutants due to the decrease in guanine concentration levels, resulting in a relative increase in hypoxanthine concentration^23^ - effectively shortening the crystal’s b-axis.

### Zebrafish crystal shape is directed by bond type, number, and interaction strength between purines

Our quantitative results on crystal morphologies using the Pnp4a-/- mutants further reinforce the recently observed connections reporting that molecular crystal morphology depends on crystal purine composition^2,30^, specifically when considering the relative purine concentrations within the crystal vesicles^23^. Hence, we set out to gain a deeper mechanistic understanding of this relationship by performing molecular simulations of crystal growth. By employing computational quantum chemistry methods and a Monte-Carlo framework, *Crystal Grower*^32,33^ (see Methods), we tested how molecular interactions, particularly interactions mimicking hypoxanthine doping in a solid solution scenario, could mechanistically influence zebrafish crystal morphology.

We began by examining the molecular structures of guanine and hypoxanthine, which differ only by the presence of a single amino (-NH2) group, which is absent in hypoxanthine molecules (Fig. 3A). Then, we analyzed the recently refined crystal structure of pure beta-guanine^17^ by computing the adjacency matrix and determining the types of intermolecular interactions using the *ToposPro*^34^ module AutoCN. This network analysis of the guanine crystal lattice identified that this -NH2 group is highly interactive with neighboring molecules, forming two hydrogen bonds (Fig. 3B). The group forms an additional five Van der Waals interactions, which display weak bond strengths.

**Figure 3.**
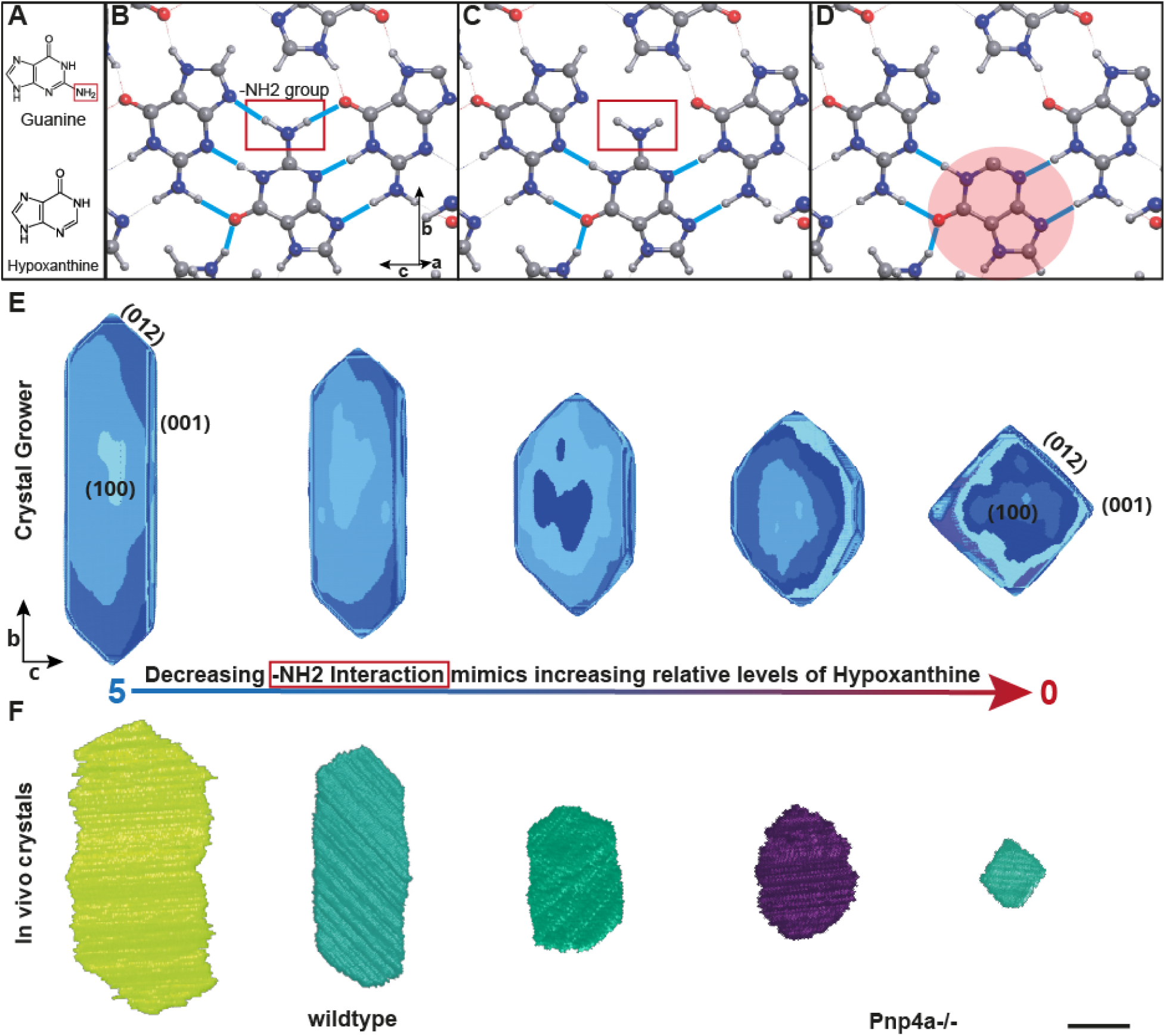
Zebrafish crystallization and crystal macromorphology depend on bond interaction strength between purines. **A** Chemical structure of guanine (top) and hypoxanthine (bottom). The two molecules only differ by a -NH2 group (red box). **B** b-c plane of the beta-guanine crystal structure. The amino group (red box) forms two hydrogen bonds (cyan) with neighboring guanine molecules. **C** b-c plane of beta-guanine crystal structure without the identified -NH2 hydrogen interactions (red box from B) corresponds to 0 interaction strength in (E). **D** b-c plane of beta-guanine crystal structure with the central guanine molecule being exchanged by hypoxanthine. This hypothetical conformation does not have the two hydrogen bonds identified in the pure guanine lattice (red box in B). **E** From left to right, simulated crystal morphologies from C in decreasing -NH2 interaction strength order (range 0-5). Note the striking similarity of simulated crystals with zero interaction strength with *in vivo* Pnp4a-/- crystals. **F** From left to right, cryoFIB-SEM 3D reconstructed crystals aligned with the closest resembling simulated crystal morphology. Scale bar: 1μm.

Considering our findings *in vivo,* where a decrease in guanine concentration leads to a relative increase in hypoxanthine concentration, but the resulting crystal lattice doping does not significantly influence crystal thickness (Fig. 2K), we focused on understanding the morphology of the (100) facet, i.e. the stereotypic hexagons present in wildtype zebrafish. Given that the guanine’s -NH2 group aligns in its orientation with the crystal’s b-axis, which is underexpressed in Pnp4a-/- mutants (Fig. 3B), we hypothesized that a higher ratio of incorporated hypoxanthine in the crystal lattice (Fig.3C) corresponds to a lower attachment probability on the (012) facet, resulting in the underexpression of the (001) facet. To test this hypothesis, we started by calculating the Voronoi partition of the guanine unit cell with ToposPro, which yields 12 generalized interactions for each molecule (see Methods). In the *Crystal Grower* framework, each molecule is represented by a proxy atom sitting at its geometric center (Fig. S2), with each generalized interaction passing through one face of the Voronoi polyhedron. We then conducted a parameter search of the 12 identified interactions to determine a parameter space of interaction scalings that resembles the hexagonal crystal morphology found in wildtype zebrafish. For simplicity, all crystals were grown under vacuum conditions without considering the energetic cost of desolvation. Remarkably, despite the clear solvent differences to *in vivo* crystallization settings, we were able to identify a parameter space (see Table S1) that produces crystals with striking morphological similarity concerning the (100) facet (Fig. 3E, left).

By identifying the generalized interactions that correspond to the hydrogen bonds between the amino group and neighboring guanine molecules (see Table S1), we tested the influence of the identified interaction (Fig. 3A-B) on the resulting crystal morphology. Interestingly, reducing this interaction strength while keeping all other interactions unchanged (Fig. 3C) leads to the (001) facet being increasingly underexpressed, i.e. shorter, similar to the square morphology of the crystals present in Pnp4a-/- mutants (Fig. 3E-F). Therefore, we conclude that this -NH2 group plays a pivotal role in determining the growth rate of the (012) facet and, consequently, the expression of the (001) facet. From a biological perspective, decreasing the absolute level of guanine molecules present in the crystal lattice in Pnp4a-/- mutants, leads to a consequent increase in the relative concentration of hypoxanthine in the iridosome (i.e. doping). Consequently, the growth rate of the (012) crystal facet is decreased, as theoretically hypothesized recently^30^. In effect, the simple presence of -NH2 interactions between guanines versus those of other purines (e.g. hypoxanthine) determines the crystal b-axis length, explaining the phenotypes obtained *in vivo* with Pnp4a-/- mutants.

## Discussion

### Zebrafish Crystal Growth is Uncoupled across Crystallographic Axes

Our study presents the first *in vivo* comprehensive analysis of the morphological development of intracellular organic crystals, using zebrafish larval iridophores as a model system. On the one hand, confocal live imaging of reflection identified an active crystal growth phase occurring during iridophore development, where exponential crystal growth and cytosolic packing occur, making these cells functional (Fig. 1A-E). On the other hand, this approach allowed us to track the growth of the (100) crystal facet from 48 to 72 hpf. The developed segmentation pipelines enabled precise extraction of 2D and 3D morphological features of single crystals, revealing significant crystal growth along the b-axis, which increased approximately four times faster than the c-axis (Fig.1H,I). This differential growth results in the crystals’ characteristic hexagonal, elongated shape. Notably, the aspect ratio between the b-axis and c-axis does not change significantly after 56 hpf, indicating that from this developmental time onwards, the final proportions of the crystals are set, and crystals increase their size while maintaining their axial proportions (Fig.1G). This points toward tight biological control across crystal shapes, which appears essential in making such crystals functional.

Given the recent reports linking crystal composition and shape^23,30^, we explored this connection further in a quantitative mechanistic manner, using a zebrafish line with impaired guanine production, *Pnp4a^cbg20^*. Crystals from homozygous Pnp4a mutants display a distinct square morphology with a significantly reduced b-axis length and an almost unaltered c-axis (Fig. 2E,F), confirming previous findings that higher relative levels of hypoxanthine hinder the (012) facet growth^23^. Quantitative live imaging of reflection coupled with cryoFIB-SEM analyses revealed that Pnp4a-/- mutants produce fewer and smaller crystals, supporting the hypothesis that the molecular composition within the iridosome influences crystal morphology. While the low crystal area and numbers in Pnp4a-/- iridophores have a profound effect on the reflective properties of these cells^23^, further investigations probing the efficiency of square versus hexagon crystal morphologies will be key to address how form directs subcellular function in this context.

Our findings indicate that the observed hexagonal morphology of the (100) facet (i.e. b- and c-axis lengths) results from the inherent geometry of the underlying H-bond network. Nevertheless, when comparing c-axes between wildtype and Pnp4a-/- crystals, we do not observe significant length differences (Fig. 2M), suggesting that incorporation of hypoxanthine in the mutant scenario does not significantly alter the purine binding probability into the (001) facet. Rather, Pnp4a-/- crystals have a disrupted H-bond network geometry due to the missing amino group, disfavouring the (012) facet but leaving the (001) facet unperturbed.

Concerning the regulation of crystal thickness, i.e. the a-axis, our work indicates that its length is also independently controlled from the other crystallographic axes, as recently suggested by colleagues^2,30^. Importantly, we found no significant differences in crystal thickness between wildtype and Pnp4a-/- iridophores. Tight control of crystal thickness is important in biological systems, as ordering and optical thickness of layers set the efficiency of multilayer reflectors^4,35^. This might underlie the differences observed between the average crystal thickness in our dataset from previously reported values in adult zebrafish skin iridosomes^29^. The mechanism by which zebrafish and other animals exert control over the thermodynamically favored π − π stacking direction has so far remained elusive. We deem a stabilization mechanism of crystal thickness based on purine doping unlikely, considering our *in vivo* analysis of Pnp4a-/- cryoFIB-SEM data (Fig. 2K). This idea has also been supported recently by *in vitro* crystallization and molecular simulation studies^30^. In addition, other reports have hypothesized that hydrophobic polymers could stabilize the thickness of the a-axis by preferential absorption on the (100) facet^36,37^, or the involvement of membrane-bound templating sheets or fibrils^4,22^.

### Crystal Composition Determines b-Axis Length *in vivo*

Our simulations of crystal growth using the Monte-Carlo framework *Crystal Grower* provided valuable insights into the sub-molecular interactions shaping crystal morphology. The presence of an amino group in guanine, absent in hypoxanthine, was found to form multiple hydrogen bonds, influencing the growth rate of the (012) facet (Fig. 3A-B). Reducing the interaction strength of this -NH2 group, simulated an increased doping of hypoxanthine in the beta-guanine crystal structure, which proved sufficient for the underexpression of the (001) facet. In line with our *in vivo* results, recent theoretical work supported by *in vitro* crystallization studies reached similar conclusions^30^. Remarkably, this is consistent with the morphologies observed in wildtype and Pnp4a-/- mutants (Fig. 3E-F and ^23^), despite ignoring solvent contributions in our crystal growth simulations. This suggests that the morphology of the (100) facet in zebrafish purine crystals might only be weakly influenced by solvent conditions.

Altogether, this leads to a unified mechanistic idea that the length of the b-crystallographic axis in biogenic crystals is mainly determined by the molecular interactions established between the purines composing the crystal lattice. Such composition appears controlled by the relative concentration of a variety of crystallized purines inside the iridosome.

How cells control diverse iridosomal purine concentrations to achieve controlled intracellular biocrystallization is an open avenue in the field. This regulation, likely dynamic, can be done at the level of intracellular transport, through respective transporter binding affinities^38,39^. Cell type-specific, dynamic purine regulation can also be achieved at a higher level by altering particular expression levels of different purine phosphorylases^23,25,40^ - effectively changing purine concentration availability in the cell. Further exploration into the cellular and molecular dynamics underlying organic crystallization across species will provide a general framework to understand the making of functional intracellular crystals.

### Limitations of the study

Here, we present comprehensive quantitative mechanistic insight into the growth and formation of zebrafish purine crystals. Despite this, reflection datasets at 72 hpf may be limited by overcrowding of crystals within iridophores, making the segmentation of single crystals difficult, hence limiting the detection of smaller crystals and rare crystal shapes. Given the nature of Pnp4a-/- mutants, with low crystal count and density, the corresponding cryoFIB-SEM datasets suffer from low numbers of measured crystals. This may explain the small changes in volume compared to the wildtype datasets. Finally, all the unit cell calculations used in simulations were run with the pure beta-guanine structure^17^. Hypoxanthine doping was only explored with regard to the missing -NH2 group, under the premise that the incorporation of guanine and hypoxanthine into the crystal lattice is the same. Given the similarity between the two molecules, and the X-ray diffraction findings that incorporation of hypoxanthine only distorts the crystal lattice in the a-axis direction^2^, we believe this assumption is reasonable.

## Methods

### Ethics Statement

This study followed European Union directives (2010/63/EU) and German law, with license #TVV52/2021 - ‘Generierung von Zebrafischlinien zur Untersuchung der Größe und Form von Organen und Organellen‘. Genetic engineering work was carried out in an S1 area (MPI-CBG, S1-Labore 4., Az.: 54-8451/103, project leader Rita Mateus), following guidelines according to Section 21, Paragraph 1 of the German Genetic Engineering Act, and within projects 01 and 03 from the Mateus laboratory, according to Section 28 of the German Genetic Engineering Safety Ordinance (GenTSV).

### Zebrafish lines and maintenance

All zebrafish (*Danio rerio*) lines were maintained in a recirculating system with a 14 h/day, 10 h/night cycle at 28°C. Crosses were performed with 3- to 12-month-old adults. Embryos were kept in E3 zebrafish embryo medium at 28.5°C until the desired developmental stage was reached. Some fish were kept at 30°C to reach the 56 hpf stage at 52 hpf. The transgenic and mutant lines used were: Et(Ola.Edar:GAL4,14xUAS:GFP)^TDL358Et^ (RRID: ZDB-FISH-150901-2380)^26^, denominated TDL358:GFP throughout the paper for simplicity, Tg(pnp4a:PALM-mCherry)^wprt10Tg^ (RRID: ZDB-FISH-210414-18)^31^, and *pnp4a^cbg20^* ^(23)^. All lines were kept in the wildtype AB strain background.

### Genotyping and phenotyping *pnp4a^cbg20^* mutants

The *pnp4a^cbg20^* mutation was kept in heterozygosity owing to this genetic alteration being lethal from juvenile stages onwards (approximately from 2 weeks old), when kept in homozygosity. As such, individual imaged larvae and breeding adults were genotyped, to confirm mutation zygosity. Briefly, genomic DNA from individual larvae or fin-clipped tail fins were extracted using the Kapa Express Extract kit (Kapa Biosystems) according to the manufacturer’s protocol. This was followed by performing PCR with KAPA2G Robust HotStart ReadyMix (Kapa Biosystems) with primers surrounding the pnp4a mutation region (FWD, 5′-CAGAATTTGTGCTTGTGTTC-3′; REV, 5′-CCTTGTACTGGTGATTGTAATG-3′). As these are located in intron regions, amplicons were specific to the mutation and not the transgenes present. Then, PCR products were sent for sequencing. Sequences were analyzed using Snapgene. Owing to the very clear phenotypes in homozygous larvae (lack of eye iridescence at 3 days post fertilization), this mutation can also be reliably identified in individual fish at this developmental stage.

### Confocal live imaging of reflection

Live imaging by point-scanning confocal microscopy was performed in larvae anesthetized in 0.1% MS-222 (Sigma) diluted in E3 embryo medium, and then mounted laterally on a concave slide (Sigma-Aldrich, BR475505) with 0.5% low-melting-point agarose (Sigma A9414) in E3 medium. This maximized the position of one of the larval eyes to be perpendicular to the incident laser. A Zeiss LSM880 Airy upright confocal microscope with a dipping W Plan-Apochromat 40x/1NA water immersion objective was used for all live imaging acquisition. Reflection imaging was performed by taking advantage of the optical properties of the flat zebrafish crystals, illuminating the larvae with a 633 nm incident laser, and adjusting the imaging detector to capture reflection arising from the crystals +/- 10nm from the incident laser line. Fluorescence and reflection imaging were acquired sequentially for each sample frame. Live imaging was repeated at least 3 times with different biological replicates per condition and zebrafish line.

### Iridophore area and reflection quantification

After full eye confocal image acquisition, maximum intensity z-projections (MIPs) of all acquired signals were obtained using Fiji^41^. The obtained images from the green fluorescent protein channel, corresponding to cytoplasmic signal from the TDL358:GFP+ cells (iridophores), were segmented using Ilastik^42^, using the pixel classification method and batch mode after initial training dataset. This provided individual binary masks that labeled the contour of all iridophores in the samples. The mask was used to define the regions of interest (ROIs) within the MIPs of other obtained channels, allowing measurements of the corresponding reflectance within iridophores and iridophore area for each sample using a custom-made Fiji macro. Individual sample reflection was normalized by dividing the reflectance signal per corresponding iridophore area, allowing comparisons between samples and fitting. Plotting and fitting was performed using Graphpad Prism.

### High-Pressure Freezing of Zebrafish Larvae

Staged zebrafish larvae at 56 or 72 hpf were anesthetized and then euthanized with 1% MS-222 (Sigma). Then, heads were removed from embryos with a scalpel blade. Three heads at a time were transferred with a cut pipette tip into high-pressure freezing carriers made from aluminum (Leica Microsystems or M. Wohlwend GmbH) with 200μm depth. To preserve the tissues in cryogenic conditions, the remaining liquid was removed and replaced by 10% (or 20%) dextran40 (MW 40 000 g/mol, Roth) in 0.1M phosphate buffer at pH 7.2. The sample volume was closed by a flat carrier (B) wetted by hexadecen (Sigma). The sample was then high-pressure frozen in a Leica ICE high-pressure freezer (Leica Microsystems, Vienna) and the frozen samples were stored in liquid nitrogen until further processing (up to 1 month).

### Cryogenic Focused Ion Beam Scanning Electron Microscopy

High-pressure frozen fish heads in aluminum carriers were transferred into a pre-cooled Quorum PP3010 cryo preparation system (Quorum Technologies) and mounted on a Zeiss sample holder under liquid nitrogen. This was then transferred into the aQuilo cryo preparation chamber before transfer onto the pre-cooled Zeiss crossbeam 550 cryoFIB-SEM (Carl Zeiss Microscopy, Oberkochen), with a cryo-stage at -160°C (-180°C anti-contaminator temperature). The sample’s surface was scanned at a working distance of 5.1 mm with electrons at 2.0 kV, 50 pA, using the InLens backscattered EsB, InLens, or SESI secondary electron detection signal to determine the position of the zebrafish eyes. Subsequently, the sample was moved back into the preparation chamber and sputter-coated with Platinum at 5 mA for 60sec. Back on the cryo-stage, the sample was tilted gradually to 54°, and the surface was imaged again for orientation. A 60-80 µm wide region containing the pigmented epithelium of the eye was selected and milled with the SmartFIB software (Carl Zeiss) using the 30nA@30kV or 65nA@30kV FIB probe current. The cross-sectioned surface was then polished using a 7nA@30kV probe current to visualize the features of interest better. The cryoFIB-SEM volumes were then acquired by serial slicing (40µm wide) and either imaging with a 3nA@30kV FIB probe current or milling round with a 2.5kV, 50pA SEM beam, resulting in a slice thickness of 32 or 65nm respectively. The InLens signal alone, or backscattered InLens (EsB) signal simultaneously, was recorded with a tilt correction of 36°. The pixel size of the acquisition was 15.4 nm/px (or 30 nm/px) in x/y direction. The image store resolution was 2048 x 1536 pixels, although a reduced area was often selected for scanning, concentrating on the region of interest, speeding up scanning time and reducing beam damage. The cycle time was kept around 20 seconds by a line averaging around N = 60 for noise reduction. Stacks of roughly 500 slices were acquired where possible.

### 2D crystal segmentation, morphometric feature extraction and quantification

Using the acquired confocal microscopy reflection images, and given that the zebrafish crystals’ a-axis is well below the diffraction limit^27,29^, we decided to treat single-plane crystal images as an effective 2D system, neglecting the crystal thickness and assuming that the (100) crystal plane is parallel to the image plane. We aimed to construct a large dataset of morphological features in a time-efficient, mostly automated and reusable way. This is a challenge as crystals tend to be close to the diffraction limit in early developmental stages, and in later stages, these are very packed, leading to reflection fringe effects due to thin film interference^35^, making them hard to segment. Therefore, we developed a novel segmentation pipeline, where we first pick regions of interest (ROIs), where crystals are well separated and fully visible. After selecting these regions, we normalize the images to their respective 99th percentile, enhancing pixel classification’s robustness and ensuring that the classifiers remain reusable across different image datasets.

Next, using the Ilastik^42^ Pixel Classification tool, which employs semantic segmentation with a random forest classification, we classify the pixels of the training image dataset as either belonging to a crystal or the background. With the pixel mask generated from this classification, we identify single crystals using a simple thresholding method (threshold set at 0.55). A size filter is then applied to exclude all objects smaller than 200 pixels (approximately 0.5µm²). Following this, the identified objects undergo further classification into “good” and “bad” crystals using the Ilastik object classifier. This step is crucial for excluding overlapping or not fully visible crystals. Specifically, object masks are classified as bad if they are concave or possess sharp edges, as these characteristics likely indicate a cluster of crystals that have been mistakenly identified as a single object. Also, any masks colliding with the image border are excluded from further analysis. The resulting masks are used to extract morphometric features, which can be measured. This was accomplished using the regionprops_table function from the scikit-image library^43^ and the label_statistics function from the Simpleitk library^44^. This approach works well for segmenting 2D crystal images up to 56 hpf. Beyond this time point, however, the cells get overcrowded with crystals, making an object-based segmentation approach unfeasible. This is primarily due to interference effects and missing spatial separation of the crystals. We used the freehand measuring tool in Fiji^41^ to segment the 72 hpf datasets and then extracted and quantified them as above.

### 3D Segmentation, morphometric feature extraction and quantification of cryoFIB-SEM images

CryoFIB-SEM image stacks are knowingly hard to segment as charging and curtaining artifacts distort the images, and intensities fluctuate strongly^45^. Therefore, we pre-processed the image stacks as previously proposed^45^, before segmenting them with a custom-trained Universal Network (U-Net). All steps to prepare and segment the images were done with *Dragonfly*. Further, the milling process between consecutive images can often lead to misalignment in the z-direction. To address this issue, we employed the Sum Squared Difference (SSD) method and manual alignment to ensure proper Z-Stack alignment. We applied a 3x3 median filter across all slices to reduce noise. Next, to counteract fluctuating intensities, we enhanced the contrast of the images by applying CLAHE (Contrast Limited Adaptive Histogram Equalization)^46^. We used a kernel size of 150 and 200 bins each, ensuring the contrast enhancement was consistent across the entire dataset. Finally, to eliminate curtaining artifacts that can arise during imaging, we utilized Dragonfly’s decurtaining tool. We hand-segmented several ROIs into a “Background” and a “Crystal” category to generate ground truth annotations. These ground truth annotations were then used to train a fully connected convolutional neural network (U-Net) inside Dragonfly for automated segmentation of FIB-SEM image stacks. Segmentation masks were corrected manually to separate crystal objects if needed. Once segmentation was complete, we extracted the crystallographic axes (a, b and c) lengths by fitting an ellipsoid to each crystal object using custom-made Python code.

### Statistical analysis

Statistical comparisons of morphological features were done using the Python library SciPy^47^ by performing a non-parametric, unpaired, two-tailed Mann-Whitney t-test (stats.ttest_ind) between the different conditions. Statistical experimental details can be found in respective figure legends. All plots were created using the Python library Seaborn^48^.

### Monte Carlo simulations using Crystal Grower

To predict crystal growth with *Crystal Grower*^32,33^, we had to prepare the raw crystal structure file. This entailed dividing the structure into Voronoi polyhedra, which are then used as minimal building blocks to reconstruct the crystal via a Monte Carlo algorithm. For this, we selected a primitive unit cell that contains the molecules of interest. We used the β-polymorph crystal structure for our simulations, which was recently refined by 3D electron diffraction^17^. Next, we calculated the adjacency matrix for the chosen structure using the AutoCN module in *ToposPro*. This step determined the types of interactions between the molecules, namely hydrogen bonds and Van der Waals interactions. Following this, we calculate the Voronoi partitioning for the molecular crystal using the *ToposPro* module *ADS*. Based on the previously obtained adjacency matrix, this calculation involves creating a proxy atom at the centroid of the molecules and combining the intermolecular interactions into generalized interactions between these proxy atoms. In our case, this results in 12 interactions, with each interaction corresponding to one side of a 12-sided Voronoi polyhedron. Finally, the obtained simplified representation is fed into the *Crystal Grower* interface, where it can be used to reconstruct the crystal via Monte Carlo Simulation. For all simulations, we used an initial driving force of Δμ = 100 kJ/mol at a temperature of T=28°C. This high driving force assures fast nucleation and crystal growth. All crystals were grown for 250000 (∼50000 particles) steps.

## DATA AND CODE AVAILABILITY

Source data for Figs. 1–3, Figs. S1-S2, and Table 1 are available in the online version of the paper. Custom code to analyze images is available upon request.

## KEY RESOURCES TABLE

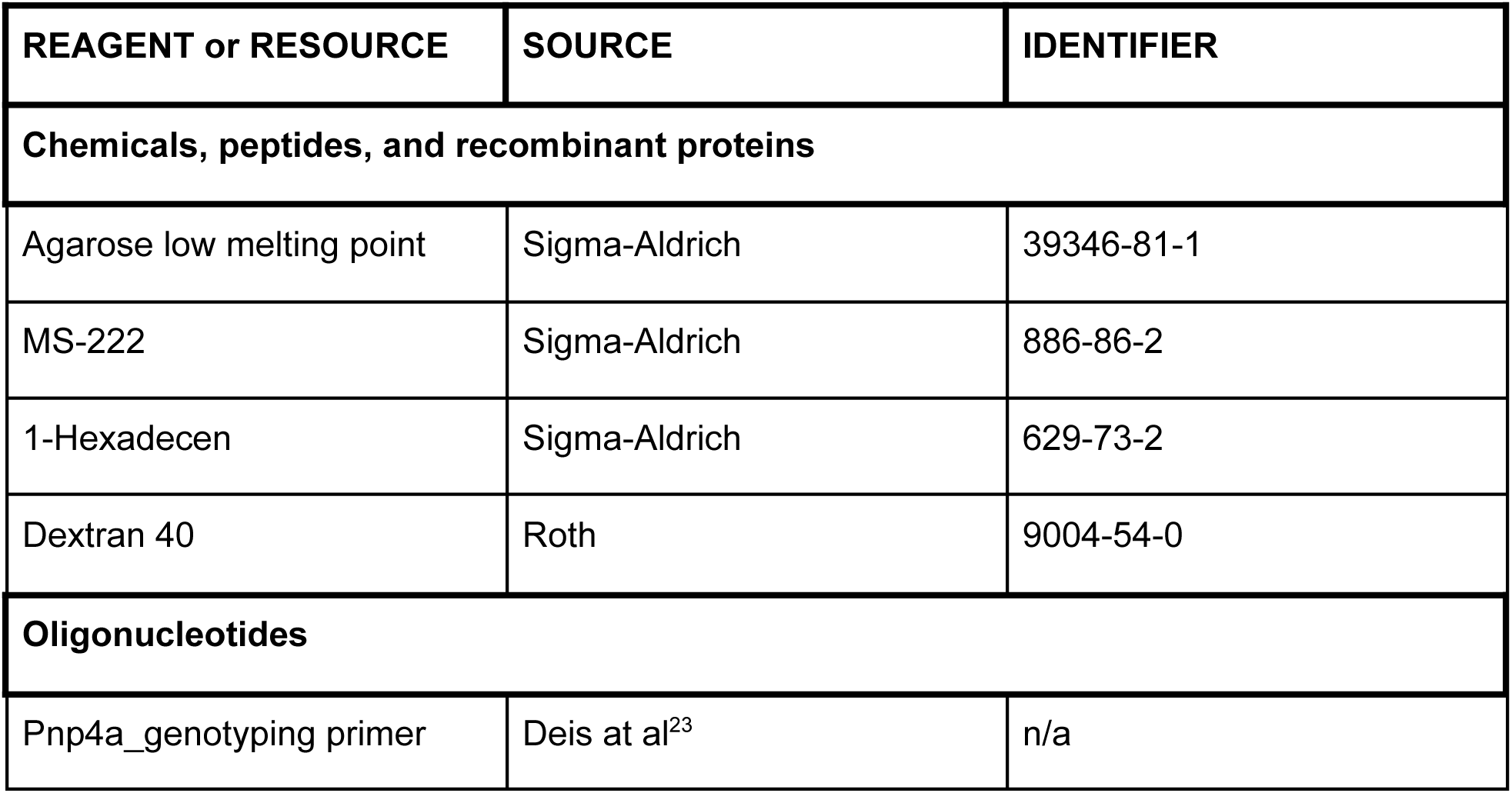
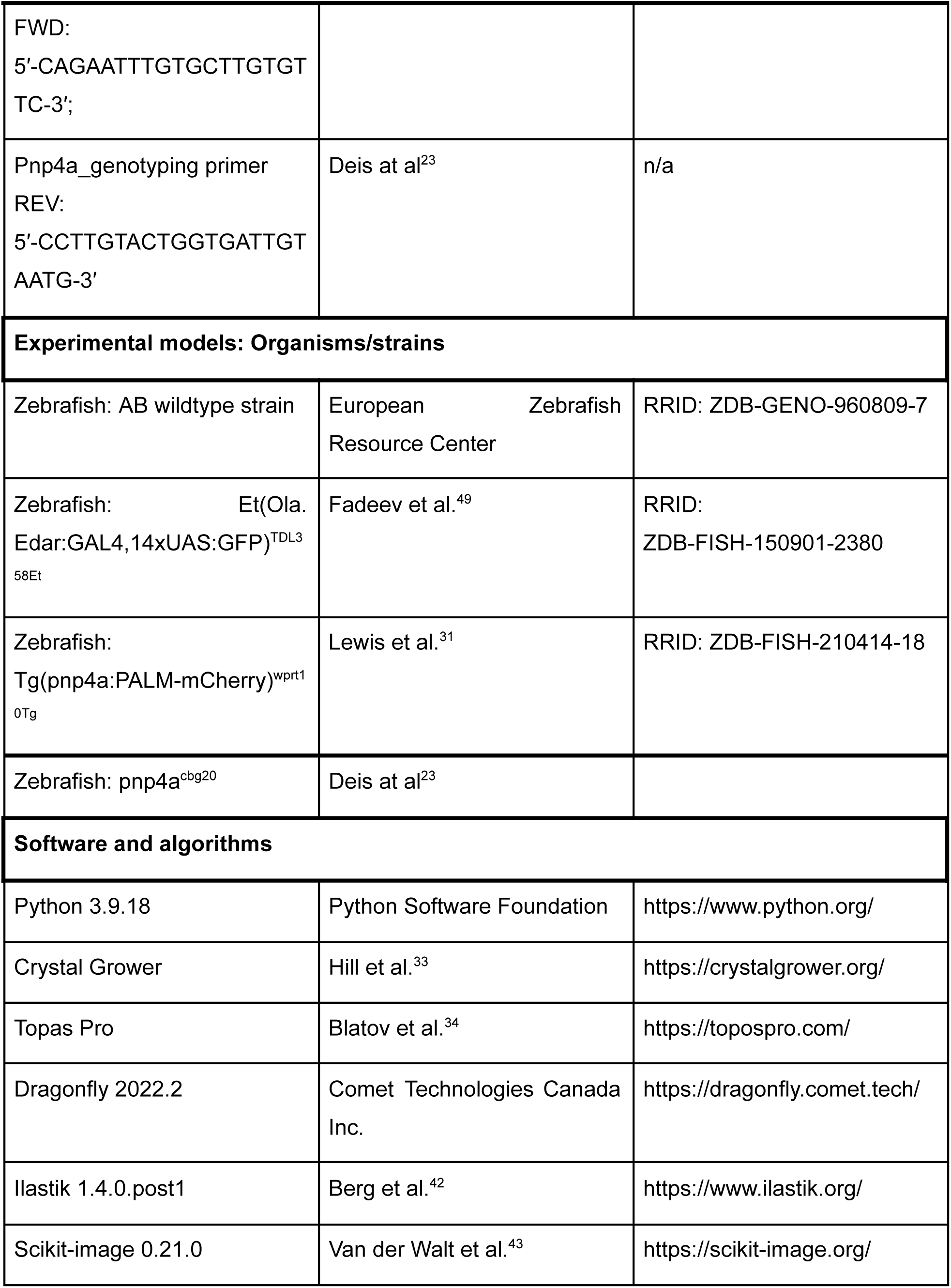
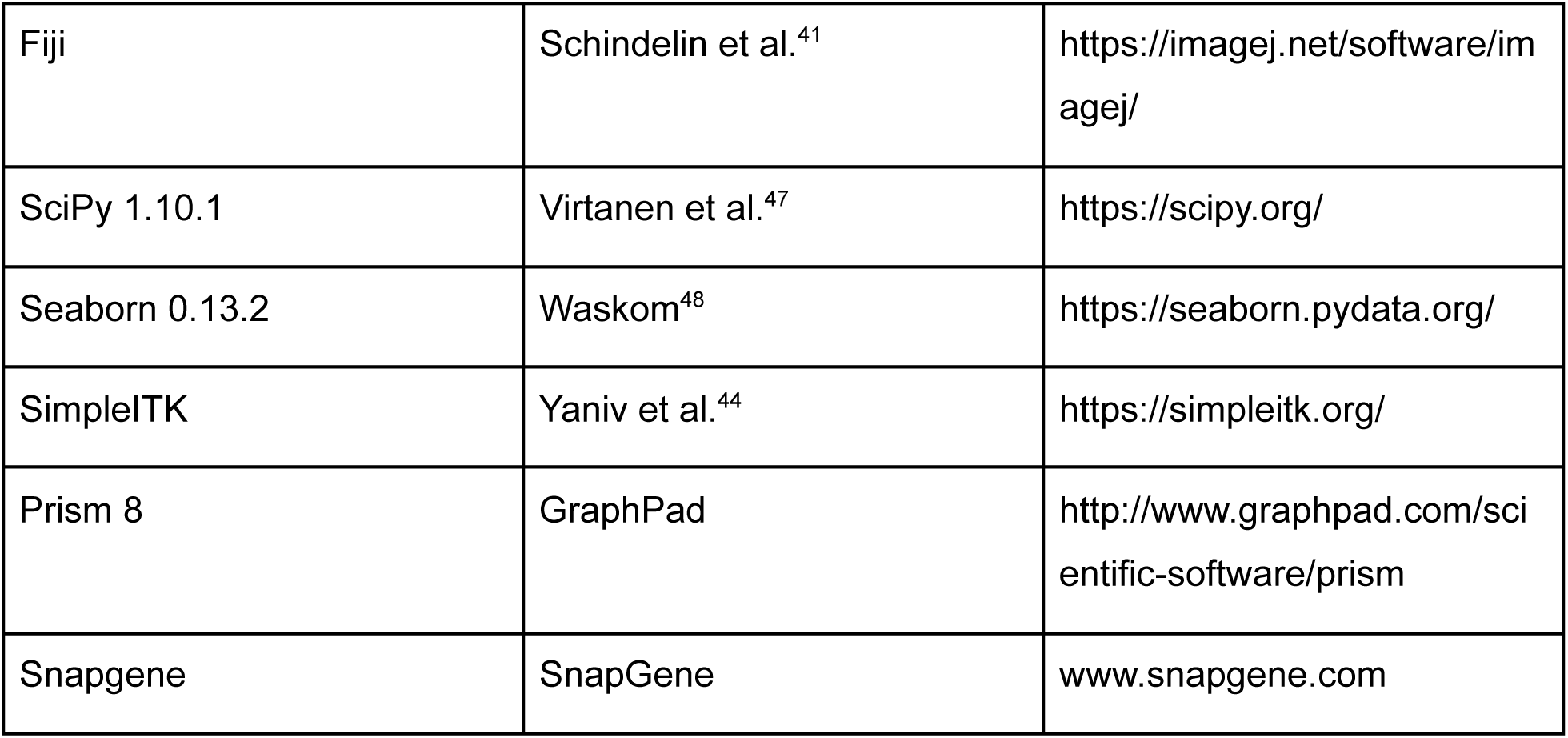

## Acknowledgements

We are grateful to Yael Politi and Luca Bertinetti for support, training, and use of the cryoFIB-SEM microscope. We thank Diego Mauro, Rajalakshmi Palaniappan, Mara Lampert, and Robert Haase for initial efforts on performing 2D crystal segmentation, the MPI-CBG Light Microscopy Facility for insightful discussions and training, and the MPI-CBG Zebrafish Team Unit for fish care. J.R., S.K., and R.M. acknowledge funding from the Max Planck Society and the Deutsche Forschungsgemeinschaft (DFG, German Research Foundation) under Germany’s Excellence 381 Strategy - EXC-2068-390729961 - Cluster of Excellence Physics of Life of TU Dresden.

## Author contributions

J.R. performed most live imaging experiments and data analysis, with the help of S.K. and R.M.; M.W-B processed samples for cryoFIB-SEM and performed respective imaging; J.R. wrote code and established the 2D and 3D data analysis pipelines as well as being the lead in performing all Monte Carlo simulations; J.R. and R.M. conceived the project and prepared the manuscript. All authors read and approved the final version of the manuscript.

## References

1. Pilátová, J., Pánek, T., Oborník, M., Čepička, I., and Mojzeš, P. (2022). Revisiting biocrystallization: purine crystalline inclusions are widespread in eukaryotes. ISME J 16, 2290–2294. 10.1038/s41396-022-01264-1.

2. Pinsk, N., Wagner, A., Cohen, L., Smalley, C.J.H., Hughes, C.E., Zhang, G., Pavan, M.J., Casati, N., Jantschke, A., Goobes, G., et al. (2022). Biogenic Guanine Crystals Are Solid Solutions of Guanine and Other Purine Metabolites. J. Am. Chem. Soc. 144, 5180–5189. 10.1021/jacs.2c00724.

3. Pavan, M.E., Movilla, F., Pavan, E.E., Di Salvo, F., López, N.I., and Pettinari, M.J. (2023). Guanine crystal formation by bacteria. BMC Biology 21, 66. 10.1186/s12915-023-01572-8.

4. Gur, D., Palmer, B.A., Weiner, S., and Addadi, L. (2017). Light Manipulation by Guanine Crystals in Organisms: Biogenic Scatterers, Mirrors, Multilayer Reflectors and Photonic Crystals. Adv Funct Materials 27, 1603514. 10.1002/adfm.201603514.

5. Teyssier, J., Saenko, S.V., Van Der Marel, D., and Milinkovitch, M.C. (2015). Photonic crystals cause active colour change in chameleons. Nat Commun 6, 6368. 10.1038/ncomms7368.

6. Gur, D., Palmer, B.A., Leshem, B., Oron, D., Fratzl, P., Weiner, S., and Addadi, L. (2015). The Mechanism of Color Change in the Neon Tetra Fish: a Light-Induced Tunable Photonic Crystal Array. Angew Chem Int Ed 54, 12426–12430. 10.1002/anie.201502268.

7. Levy-Lior, A., Shimoni, E., Schwartz, O., Gavish-Regev, E., Oron, D., Oxford, G., Weiner, S., and Addadi, L. (2010). Guanine-Based Biogenic Photonic-Crystal Arrays in Fish and Spiders. Adv Funct Materials 20, 320–329. 10.1002/adfm.200901437.

8. Kobelt, F., and Linsenmair, K.E. (1992). Adaptations of the reed frog Hyperolius viridiflavus (Amphibia: Anura: Hyperoliidae) to its arid environment: VI. The iridophores in the skin as radiation reflectors. J Comp Physiol B 162. 10.1007/BF00260758.

9. Retinal structure in Latimeria chalumnae (1973). Phil. Trans. R. Soc. Lond. B 266, 493–518. 10.1098/rstb.1973.0054.

10. Palmer, B.A., Taylor, G.J., Brumfeld, V., Gur, D., Shemesh, M., Elad, N., Osherov, A., Oron, D., Weiner, S., and Addadi, L. (2017). The image-forming mirror in the eye of the scallop. Science 358, 1172–1175. 10.1126/science.aam9506.

11. Jordan, T.M., Partridge, J.C., and Roberts, N.W. (2012). Non-polarizing broadband multilayer reflectors in fish. Nature Photon 6, 759–763. 10.1038/nphoton.2012.260.

12. Chandel, N.S. (2021). Nucleotide Metabolism. Cold Spring Harb Perspect Biol 13, a040592. 10.1101/cshperspect.a040592.

13. Sivaguru, M., Saw, J.J., Wilson, E.M., Lieske, J.C., Krambeck, A.E., Williams, J.C., Romero, M.F., Fouke, K.W., Curtis, M.W., Kear-Scott, J.L., et al. (2021). Human kidney stones: a natural record of universal biomineralization. Nat Rev Urol 18, 404–432. 10.1038/s41585-021-00469-x.

14. Mei, Y., Dong, B., Geng, Z., and Xu, L. (2022). Excess Uric Acid Induces Gouty Nephropathy Through Crystal Formation: A Review of Recent Insights. Frontiers in Endocrinology 13.

15. Petratou, K., Subkhankulova, T., Lister, J.A., Rocco, A., Schwetlick, H., and Kelsh, R.N. (2018). A systems biology approach uncovers the core gene regulatory network governing iridophore fate choice from the neural crest. PLoS Genet 14, e1007402. 10.1371/journal.pgen.1007402.

16. Miyadai, M., Takada, H., Shiraishi, A., Kimura, T., Watakabe, I., Kobayashi, H., Nagao, Y., Naruse, K., Higashijima, S., Shimizu, T., et al. (2023). A gene regulatory network combining Pax3/7, Sox10 and Mitf generates diverse pigment cell types in medaka and zebrafish. Development 150, dev202114. 10.1242/dev.202114.

17. Wagner, A., Merkelbach, J., Samperisi, L., Pinsk, N., Kariuki, B.M., Hughes, C.E., Harris, K.D.M., and Palmer, B.A. (2024). Structure Determination of Biogenic Crystals Directly from 3D Electron Diffraction Data. Crystal Growth & Design, acs.cgd.3c01290. 10.1021/acs.cgd.3c01290.

18. Wagner, A., Ezersky, V., Maria, R., Upcher, A., Lemcoff, T., Aflalo, E.D., Lubin, Y., and Palmer, B.A. (2022). The Non-Classical Crystallization Mechanism of a Composite Biogenic Guanine Crystal. Advanced Materials 34, 2202242. 10.1002/adma.202202242.

19. Kelsh, R.N., Brand, M., Jiang, Y.-J., Heisenberg, C.-P., Lin, S., Haffter, P., Odenthal, J., Mullins, M.C., Eeden, F.J.M.V., Furutani-Seiki, M., et al. (1996). Zebrafish pigmentation mutations and the processes of neural crest development. Development 123, 369–389. 10.1242/dev.123.1.369.

20. Hirata, M., Nakamura, K.I., Kanemaru, T., Shibata, Y., and Kondo, S. (2003). Pigment cell organization in the hypodermis of zebrafish. Developmental Dynamics 227, 497–503. 10.1002/dvdy.10334.

21. Hirata, M., Nakamura, K.I., and Kondo, S. (2005). Pigment cell distributions in different tissues of the zebrafish, with special reference to the striped pigment pattern. Developmental Dynamics 234, 293–300. 10.1002/dvdy.20513.

22. Eyal, Z., Deis, R., Varsano, N., Dezorella, N., Rechav, K., Houben, L., and Gur, D. (2022). Plate-like Guanine Biocrystals Form via Templated Nucleation of Crystal Leaflets on Preassembled Scaffolds. J. Am. Chem. Soc. 144, 22440–22445. 10.1021/jacs.2c11136.

23. Deis, R., Lerer-Goldshtein, T., Baiko, O., Eyal, Z., Brenman-Begin, D., Goldsmith, M., Kaufmann, S., Heinig, U., Dong, Y., Lushchekina, S., et al. (2024). Genetic control over biogenic crystal morphogenesis in zebrafish. Nat Chem Biol. 10.1038/s41589-024-01722-1.

24. Kimura, T., Takehana, Y., and Naruse, K. (2017). Pnp4a Is the causal gene of the medaka iridophore Mutant guanineless. G3: Genes, Genomes, Genetics 7, 1357–1363. 10.1534/g3.117.040675.

25. Pedley, A.M., and Benkovic, S.J. (2017). A New View into the Regulation of Purine Metabolism: The Purinosome. Trends in Biochemical Sciences 42, 141–154. 10.1016/j.tibs.2016.09.009.

26. Levesque, M.P., Krauss, J., Koehler, C., Boden, C., and Harris, M.P. (2013). New tools for the identification of developmentally regulated enhancer regions in embryonic and adult zebrafish. Zebrafish 10, 21–29. 10.1089/zeb.2012.0775.

27. Webb, R.H. (1996). Confocal optical microscopy. Rep. Prog. Phys. 59, 427–471. 10.1088/0034-4885/59/3/003.

28. Schertel, A., Snaidero, N., Han, H.-M., Ruhwedel, T., Laue, M., Grabenbauer, M., and Möbius, W. (2013). Cryo FIB-SEM: Volume imaging of cellular ultrastructure in native frozen specimens. Journal of Structural Biology 184, 355–360. 10.1016/j.jsb.2013.09.024.

29. Gur, D., Bain, E.J., Johnson, K.R., Aman, A.J., Pasolli, H.A., Flynn, J.D., Allen, M.C., Deheyn, D.D., Lee, J.C., Lippincott-Schwartz, J., et al. (2020). In situ differentiation of iridophore crystallotypes underlies zebrafish stripe patterning. Nat Commun 11, 6391. 10.1038/s41467-020-20088-1.

30. Wagner, A., Hill, A., Lemcoff, T., Livne, E., Avtalion, N., Casati, N., Kariuki, B.M., Graber, E.R., Harris, K.D.M., Cruz-Cabeza, A.J., et al. (2024). Rationalizing the Influence of Small-Molecule Dopants on Guanine Crystal Morphology. Chem. Mater., acs.chemmater.4c01771. 10.1021/acs.chemmater.4c01771.

31. Lewis, V.M., Saunders, L.M., Larson, T.A., Bain, E.J., Sturiale, S.L., Gur, D., Chowdhury, S., Flynn, J.D., Allen, M.C., Deheyn, D.D., et al. (2019). Fate plasticity and reprogramming in genetically distinct populations of *Danio* leucophores. Proc. Natl. Acad. Sci. U.S.A. 116, 11806–11811. 10.1073/pnas.1901021116.

32. Anderson, M.W., Gebbie-Rayet, J.T., Hill, A.R., Farida, N., Attfield, M.P., Cubillas, P., Blatov, V.A., Proserpio, D.M., Akporiaye, D., Arstad, B., et al. (2017). Predicting crystal growth via a unified kinetic three-dimensional partition model. Nature 544, 456–459. 10.1038/nature21684.

33. Hill, A.R., Cubillas, P., Gebbie-Rayet, J.T., Trueman, M., De Bruyn, N., Harthi, Z.A., Pooley, R.J.S., Attfield, M.P., Blatov, V.A., Proserpio, D.M., et al. (2021). *CrystalGrower* : a generic computer program for Monte Carlo modelling of crystal growth. Chem. Sci. 12, 1126–1146. 10.1039/D0SC05017B.

34. Blatov, V.A., Shevchenko, A.P., and Proserpio, D.M. (2014). Applied Topological Analysis of Crystal Structures with the Program Package ToposPro. Crystal Growth & Design 14, 3576–3586. 10.1021/cg500498k.

35. Land, M.F. (1972). The physics and biology of animal reflectors. Progress in Biophysics and Molecular Biology 24, 75–106. 10.1016/0079-6107(72)90004-1.

36. Chen, F., Liu, Y., Li, L., Qi, L., and Ma, Y. (2020). Synthesis of Bio-Inspired Guanine Microplatelets: Morphological and Crystallographic Control. Chemistry A European J 26, 16228–16235. 10.1002/chem.202003156.

37. Chen, F., Guo, D., Gao, J., and Ma, Y. (2021). Bioinspired Crystallization of Guanine. J. Phys. Chem. Lett. 12, 11695–11702. 10.1021/acs.jpclett.1c03010.

38. Boswell-Casteel, R.C., and Hays, F.A. (2017). Equilibrative nucleoside transporters—A review. Nucleosides, Nucleotides & Nucleic Acids 36, 7–30. 10.1080/15257770.2016.1210805.

39. Young, J.D., Yao, S.Y.M., Baldwin, J.M., Cass, C.E., and Baldwin, S.A. (2013). The human concentrative and equilibrative nucleoside transporter families, SLC28 and SLC29. Molecular Aspects of Medicine 34, 529–547. 10.1016/j.mam.2012.05.007.

40. Bzowska, A., Kulikowska, E., and Shugar, D. (2000). Purine nucleoside phosphorylases: properties, functions, and clinical aspects. Pharmacology & Therapeutics 88, 349–425. 10.1016/S0163-7258(00)00097-8.

41. Schindelin, J., Arganda-Carreras, I., Frise, E., Kaynig, V., Longair, M., Pietzsch, T., Preibisch, S., Rueden, C., Saalfeld, S., Schmid, B., et al. (2012). Fiji: An open-source platform for biological-image analysis. Nature Methods 9, 676–682. 10.1038/nmeth.2019.

42. Berg, S., Kutra, D., Kroeger, T., Straehle, C.N., Kausler, B.X., Haubold, C., Schiegg, M., Ales, J., Beier, T., Rudy, M., et al. (2019). ilastik: interactive machine learning for (bio)image analysis. Nat Methods 16, 1226–1232. 10.1038/s41592-019-0582-9.

43. Van Der Walt, S., Schönberger, J.L., Nunez-Iglesias, J., Boulogne, F., Warner, J.D., Yager, N., Gouillart, E., and Yu, T. (2014). scikit-image: image processing in Python. PeerJ 2, e453. 10.7717/peerj.453.

44. Yaniv, Z., Lowekamp, B.C., Johnson, H.J., and Beare, R. (2018). SimpleITK Image-Analysis Notebooks: a Collaborative Environment for Education and Reproducible Research. J Digit Imaging 31, 290–303. 10.1007/s10278-017-0037-8.

45. Roldán, D., Redenbach, C., Schladitz, K., Kübel, C., and Schlabach, S. (2024). Image quality evaluation for FIB-SEM images. Journal of Microscopy 293, 98–117. 10.1111/jmi.13254.

46. Pizer, S.M., Amburn, E.P., Austin, J.D., Cromartie, R., Geselowitz, A., Greer, T., Ter Haar Romeny, B., Zimmerman, J.B., and Zuiderveld, K. (1987). Adaptive histogram equalization and its variations. Computer Vision, Graphics, and Image Processing 39, 355–368. 10.1016/S0734-189X(87)80186-X.

47. Virtanen, P., Gommers, R., Oliphant, T.E., Haberland, M., Reddy, T., Cournapeau, D., Burovski, E., Peterson, P., Weckesser, W., Bright, J., et al. (2020). SciPy 1.0: fundamental algorithms for scientific computing in Python. Nat Methods 17, 261–272. 10.1038/s41592-019-0686-2.

48. Waskom, M. (2021). seaborn: statistical data visualization. JOSS 6, 3021. 10.21105/joss.03021.

49. Fadeev, A., Krauss, J., Frohnhöfer, H.G., Irion, U., and Nüsslein-Volhard, C. (2015). Tight Junction Protein 1a regulates pigment cell organisation during zebrafish colour patterning. eLife 4, e06545. 10.7554/eLife.06545.

