## Supplemental Information for "The Role of Purine Interactions in Biogenic Crystal Shape Determination"

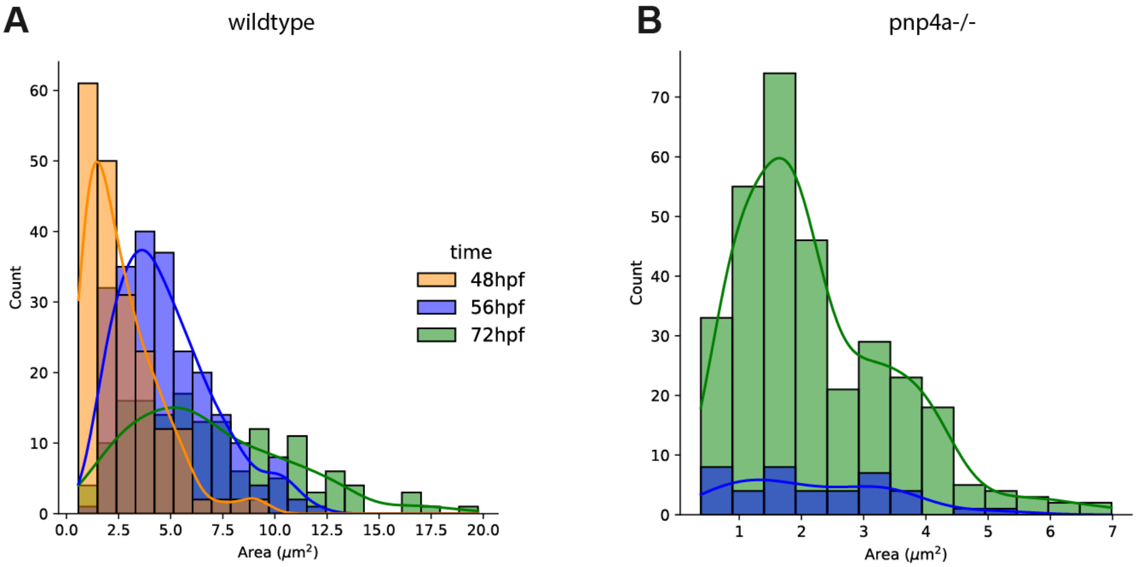


**Figure S1: Histograms of (100)-facet area of 2D segmented crystals. A-B** Crystal amount versus segmented area at 48 hpf (orange), 56 hpf (blue) and 72 hpf (green), in wildtype (**A**) and Pnp4a-/- mutants (**B**). Line shows kernel density estimation per developmental time point.

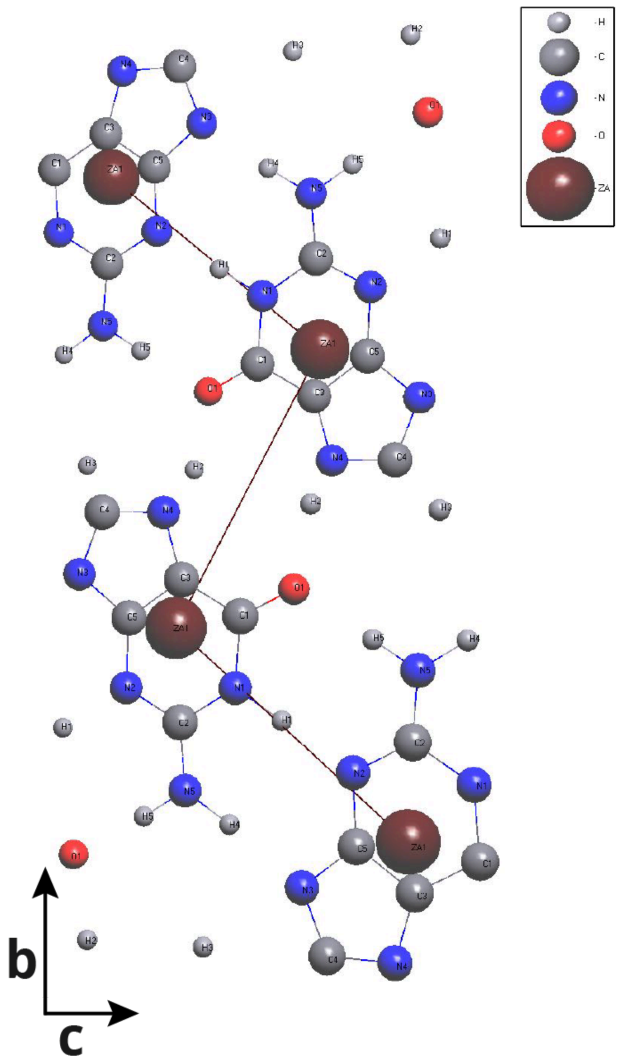


**Figure S2. Simulated b-c plane of the guanine crystal lattice.** Note that each molecule is represented with a proxy atom (see color legend, top right) at the geometric center of the guanine molecules.

| Name | Space Position | Scaling |
|---|---|---|
| A | (-1,0,0) | 1 |
| B | (1,0,0) | 1 |
| C | (x,1/2-y,1/2+z)(-1,0,0) | 1 |
| D | (x,1/2-y,1/2+z)(1,0,1) | 1 |
| E | (x,1/2-y,1/2+z)(0,0,1) | 3 |
| F | (x,1/2-y,1/2+z) | 4 |
| G | (-x,-y,-z) | 2 |
| H | (-x,-y,-z)(1,0,0) | 2 |
| I | (-x,-y,-z)(2,0,1) | 0 |
| J | (-x,-y,-z)(1,0,1) | 3 |
| K | (-x,-y,-z)(-1,-1,0) | 0 to 5 |
| L | (-x,1/2+y,1/2-z)(1,0,0) | 0 to 5 |

**Table S1. Interaction scalings from parameter search.**
Interactions in the 1st column (Name) with respective space positions of the proxy atoms, plus interaction scalings used for the simulations in Crystal Grower. Green: Van der Waals interaction responsible for pi-pi stacking. Black: Mixed interactions (VdW and H-bonding). Red: H-Bond interactions of the -NH2 residue.
